# Inter-user comparison for quantification of superparamagnetic iron oxides with magnetic particle imaging across two institutions highlights a need for standardized approaches

**DOI:** 10.1101/2023.04.03.535446

**Authors:** Hayden J. Good, Olivia C. Sehl, Julia J. Gevaert, Bo Yu, Maryam A. Berih, Sebastian A. Montero, Carlos M. Rinaldi-Ramos, Paula J. Foster

**Author notes:** Corresponding Author: Hayden J. Good. These authors contributed equally to this work (Hayden J. Good, Olivia C. Sehl).

## Abstract

**Purpose:** Magnetic particle imaging (MPI) is being explored in biological contexts that require accurate and reproducible quantification of superparamagnetic iron oxide nanoparticles (SPIONs). While many groups have focused on improving imager and SPION design to improve resolution and sensitivity, few have focused on improving quantification and reproducibility of MPI. The aim of this study was to compare MPI quantification results by two different systems and the accuracy of SPION quantification performed by multiple users at two institutions.

**Procedures:** Six users (3 from each institute) imaged a known amount of Vivotrax+ (10 μg Fe), diluted in a small (10 μL) or large (500 μL) volume. These samples were imaged with or without calibration standards in the field of view, to create a total of 72 images (6 users x triplicate samples x 2 sample volumes x 2 calibration methods). These images were analyzed by the respective user with two region of interest (ROI) selection methods. Image intensities, Vivotrax+ quantification, and ROI selection was compared across users, within and across institutions.

**Results:** MPI imagers at two different institutes produce significantly different signal intensities, that differ by over 3 times for the same concentration of Vivotrax+. Overall quantification yielded measurements that were within ± 20% from ground truth, however SPION quantification values obtained at each laboratory were significantly different. Results suggest that the use of different imagers had a stronger influence on SPION quantification compared to differences arising from user error. Lastly, calibration conducted from samples in the imaging field of view gave the same quantification results as separately imaged samples.

**Conclusions:** This study highlights that there are many factors that contribute to the accuracy and reproducibility of MPI quantification, including variation between MPI imagers and users, despite pre-defined experimental set up, image acquisition parameters, and ROI selection analysis.

## Introduction

Magnetic particle imaging (MPI) directly detects and images the spatial distribution of superparamagnetic iron oxide nanoparticles (SPIONs) with high specificity and sensitivity. Nonlinear dynamic SPION magnetization is driven using an alternating excitation magnetic field, resulting in a signal that is measured via pick up coils [1–4], with signal strength directly related to the amount of SPIONs present. Localization of SPIONs is achieved using a gradient selection field with a central region with negligible magnetic field (the field free region, FFR), compared to the magnitude of the excitation field. SPIONs outside the FFR have their magnetization saturated by the gradient selection field and thus do not respond to the excitation field. Thus, the FFR defines the region in which SPIONs can respond to the excitation field, generating a signal, and is scanned across the field of view (FOV) to produce an image. SPION tracking with MPI has numerous applications, including cell tracking [5–7], vascular/perfusion imaging [8,9], oncology imaging [10], pulmonary imaging [11,12], imaging of inflammation [13], and evaluating SPION distribution for magnetic hyperthermia [14,15]. In all these applications, accurate and reproducible quantification of SPION distribution is essential for progress in the field.

The number of preclinical commercial and prototype MPI systems is growing rapidly. There are currently two preclinical imagers available from Bruker Biospin and Magnetic Insight, Inc. Between these systems, there are differences in the configuration of the FFR, selection gradient field strength, excitation field frequency and amplitude, bore size, and available FOV [16]. The Bruker system is designed for high temporal resolution images at the cost of sensitivity with a weaker selection gradient field while the MOMENTUM system is designed for high sensitivity imaging at the cost of temporal resolution with a stronger selection gradient field [4]. Our laboratories are equipped with the Magnetic Insight, Inc., MOMENTUM^TM^ imager, hence we will focus on this type of imager in this contribution. The first MOMENTUM^TM^ system made by Magnetic Insight Inc. was installed at Stanford University in 2016 and to date there are 13 systems operating in academic and research institutions in North America, Europe, China, and Australia. This system uses a FFL (line) selection field gradient for spatial localization and has a 6 cm bore opening. While these imagers produce signal in a similar manner, each new imager differs slightly from the previous one due to continuous improvement in design and because each imager is manually calibrated. This is to be expected at this relatively early stage of MPI technology development. However, this also implies that imager performance, ostensibly in terms of resolution, sensitivity, and quantification accuracy and precision will likely vary among imagers.

To date, there has not been a comparison of the performance and resulting findings between different MOMENTUM^TM^ imagers. We expect that studies comparing imager performance would benefit the community by quantifying variation in imager performance, facilitating comparisons among laboratories and progress in the field. Furthermore, because MPI is still relatively new, the field lacks standard protocols for image acquisition and analysis, both of which can be expected to introduce potential variations in SPION quantification between imagers in different laboratories and between users in the same imager. This manuscript is focused on an inter-user study conducted with the MOMENTUM^TM^ systems located at Robarts Research Institute (RRI) and University of Florida (UF). Standardized protocols were used at each site to acquire MPI data using the same tracers by several users and the resulting data was analyzed using several approaches. As such, this is the first study to directly compare MPI quantification accuracy and precision between MPI imagers in different sites and between users within individual sites.

There are numerous SPIONs available for generating signal in MPI and each produce different signal characteristics. Ferucarbotran (now available as Vivotrax, Magnetic Insight Inc.), originally developed for magnetic resonance imaging (MRI), has been widely used in MPI work and can be considered a “gold standard” because of its ease of commercial availability even though its properties are not ideal for MPI resolution and sensitivity. Ferucarbotran has been reported to have a bimodal core size distribution; approximately 30% of the cores are 25-30 nm and 70% of the cores are ∼5 nm [17,18]. A size-fractionated product, VivoTrax+, is now available with improved MPI resolution and sensitivity performance [19]. In this study we use VivoTrax+ as a standard tracer to compare the effect of MPI imager and image analysis method on SPION quantification precision and accuracy.

Calibration curves relating known SPION mass to signal are used to quantify SPION mass distribution using MPI. While the MPI signal output is directly related to the amount of SPIONs, the output is in arbitrary units (a.u.) that are expected to vary across imagers. Calibration curves relating this imager-dependent signal strength to SPION mass are constructed for each SPION type by imaging reference phantoms (also commonly referred to as fiducials or calibration samples) which have defined, known iron mass. It is essential that these fiducials are imaged using the same parameters, including selection field gradient strength and excitation amplitude and frequency, as these factors contribute to MPI signal strength. These fiducials may be included with the subject within the imaging field of view (FOV) [20–23], which we refer to as “internal calibration”. The added benefit of internal fiducials is they can serve as benchmarks for co-registration with other imaging modalities (e.g., optical images acquired prior to a scan, computed tomography, magnetic resonance imaging, etc.). However, there is a limited number of calibration samples that can be resolved within a FOV and their signals may overlap with or interfere with signals belonging to the specimen of interest. For this reason, calibration samples may be imaged in separate image(s) [6, 10, 24, 25] and in this paper we refer to this as “external calibration”. Prior to this study it was unknown how quantification results are impacted using internal or external calibration fiducials.

The central aim of this study was to assess the reproducibility of MPI data acquisition using two different MPI imagers and of quantification of Vivotrax+ performed by six users at two academic centers. To achieve this, we compare image characteristics for the two systems and assess factors which may lead to differences in quantification including methods of calibration (internal vs. external), inter-user variability, software for analysis, and methods to select the regions-of-interest (ROI) for quantification. We aimed to develop and validate protocols that are reproducible across users and among institutions.

## Methods

Six users (3 from RRI and 3 from UF) performed the imaging experiments and analyses described below independently. One user from each institution performed additional statistical analysis and MPI relaxometry. The following aspects of this study were carefully controlled to compare user accuracy and reproducibility: sample preparation, experimental set up, image acquisition parameters, image segmentation methods, and calibration methods.

### Sample Preparation

Aliquots from the same bottle of Vivotrax+ (Magnetic Insight Inc.) were used to prepare samples by each user group. The iron concentration for the specific batch of Vivotrax+ used was measured as 4.59 mg Fe/mL, determined by the 1,10-phenanthroline assay [26]. Aliquots were stored at 4°C and prior to the experiment were brought to room temperature and vortexed to ensure a homogenous distribution of particles in solution. Samples were prepared as either calibration fiducials or test samples for quantification. The calibration fiducials were prepared with 40, 20, 10, 5, and 2.5 μg Fe in 10 μL of deionized water. Briefly, for the first sample in the dilution series, 25.3 μL of stock Vivotrax+ was mixed with 3.7 μL deionized water diluent for a final concentration of 4.0 μg Fe/μL. From this total volume, 15 μL was removed and placed in a separate container and 10 μL from this 15 μL was loaded into capillary tubing for a final sample containing 40 μg Fe in 10 μL. Then, the remaining 14.1μL were diluted further, and 10 μL placed in another fiducial. This dilution series method was used to minimize pipetting error in producing calibration standards. All subsequent samples in the dilution series were similarly prepared following the volumes outlined in Table 1. The first test sample, “Sample 1”, was prepared similarly to calibration fiducial 3, where 10 μg Fe in 10 μL deionized water was loaded into a capillary tube (n = 3 samples, shown in Figure 1a). The second test sample, “Sample 2”, was prepared by diluting 10 μg Fe in 500 μL deionized water and loaded into a PCR tube (n = 3 samples, shown in Figure 1a). All capillary tube ends were capped and sealed with Cha-Seal to prevent evaporation of samples. All images were acquired immediately after sample preparation.

**Figure 1:**
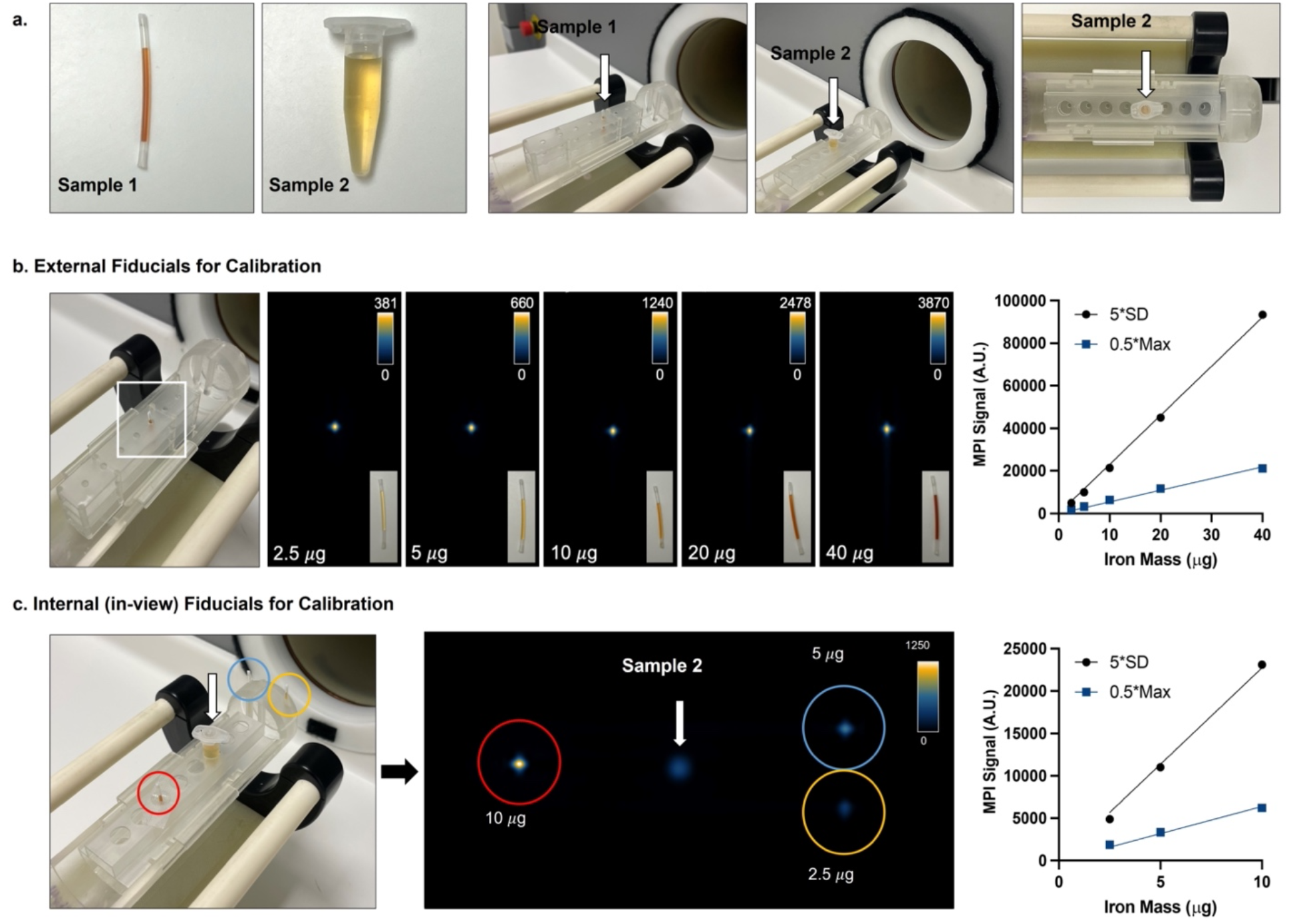
Experimental setup. (a) Two test samples were prepared, each containing 10 *μ*g of Vivotrax+ (n = 3 per user). Vivotrax+ was mixed into deionized water to a volume of 10 *μ*L (Sample 1) or 500 *μ*L (Sample 2), then loaded into capillary tubing or 0.5 mL Eppendorf tubes, respectively. Custom 3D printed holders were created to hold each tube vertically at the center of the MPI sample holder. (b) For external calibration, 5 samples of Vivotrax+ (2.5 *μ*g, 5 *μ*g, 10 *μ*g, 20 *μ*g, and 40 *μ*g, each in a total volume of 10 *μ*L) were imaged individually in capillary tubing. MPI images are shown in full dynamic range. The signal from these 5 images was quantified using 5*SD_avg_ and 0.5*Max segmentation methods, giving a linear relationship between iron mass and measured MPI signal. (c) For internal calibration, 3 fiducials containing 10 *μ*g, 5 *μ*g, and 2.5 *μ*g Vivotrax+ in total volume 10 *μ*L were placed within 3D printed scaffolds in the imaging field of view, surrounding Sample 1 or 2, as shown in the photo. These 3 calibration signals measured from each image were used to create calibration curves for both segmentation methods (5*SD_avg_ and 0.5*Max).

**Table 1:** Calibration sample preparation. All users took stock Ferucarbotran and diluted with DI water according to this table. Then, 10uL of these solutions were loaded into tubing to be imaged.

| Calibration Fiducial | Final Concentration ($\mu\text{g Fe}/\mu\text{L}$) | Solution Used | Volume of Solution ($\mu\text{L}$) | Volume of Diluent ($\mu\text{L}$) | Total Volume ($\mu\text{L}$) | Remaining Volume ($\mu\text{L}$) |
| --- | --- | --- | --- | --- | --- | --- |
| 1 | 4.0 | Stock | 25.3 | 3.7 | 29.1 | 15.00 |
| 2 | 2.0 | Calibration 1 | 14.1 | 14.1 | 28.1 | 15.00 |
| 3 | 1.0 | Calibration 2 | 13.1 | 13.1 | 26.3 | 15.00 |
| 4 | 0.50 | Calibration 3 | 11.3 | 11.3 | 22.5 | 15.00 |
| 5 | 0.25 | Calibration 4 | 7.5 | 7.5 | 15.0 | 15.00 |

### Image Acquisition

All images were acquired using the Momentum MPI field-free line imager (Magnetic Insight Inc.) either at RRI (installation date July 2019) or UF (installation date February 2019). Images were acquired in 2D with a 5.7 T/m selection field gradient and excitation field strengths of 20 and 26 mT in the X and Z channels, respectively, at RRI. Similarly, images were acquired in 2D with a 5.7 T/m selection field gradient and excitation field strengths of 16 and 19 mT in the X and Z channels, respectively, at UF. In both instruments the excitation frequency was 45 kHz. These conditions correspond to the standard imaging mode for each system, as specified by the manufacturer. These 2D images took ∼2 minutes to acquire for a 12 (Z channel) x 6 (X channel) cm FOV.

Samples were placed vertically in custom 3D printed holders (shown Figure 1a). The sample holders were tested for iron contamination by imaging with MPI to ensure there was no signal contribution from the sample holder. To measure the average background signal in the absence of an iron sample, three images of an empty sample holder were acquired. The standard deviation of the voxel values from these three images were measured and an average background signal was calculated (SDavg).

Five external calibration fiducials were individually imaged in the middle of the sample holder, central to the imaging FOV (Figure 1b). Triplicates of Sample 1 and Sample 2 were individually imaged in the center of the imaging FOV. These 6 images were repeated with 10 μg, 5 μg, and 2.5 μg calibration fiducials positioned around each test sample (layout shown in Figure 1c).

### Image Analysis

Histograms were created in MATLAB to display MPI image values for individual samples imaged at RRI and UF. MPI total signal generated by fiducials and test samples was measured in images by creating ROIs. The first ROI segmentation method set a lower threshold of five times the user’s average standard deviation of background noise (5*SDavg) corresponding to the Rose criterion [27]. For images with internal (in-view) fiducials, manual snips were performed to separate ROIs if they included overlapping signals. ROIs were analyzed again using an alternate segmentation method: delineation of the minimum threshold boundary at half the maximum signal intensity value within that ROI (0.5*Max). The maximum value for each signal was unique, therefore each ROI segmentation used a different lower threshold value which scaled with the maximum value of the ROI. MPI total signal was calculated by multiplying the mean signal of voxels in the ROI by the ROI area (mm2). Participants from RRI performed this analysis in open-source HorosTM software [28] and participants from UF conducted the analysis in open-source 3D Slicer software [29,30]. Each user analyzed their respective image data. Additionally, each user was paired with another user from the other institution to exchange and analyze each other’s images to assess differences between users and software programs.

Calibration curves map the linear relationship between iron mass of calibration samples (x-axis) and the MPI total signal generated (y-axis) to give a linear equation relating MPI total signal and iron mass. The slope of the calibration curve represents the amount of MPI signal (a.u.) generated per iron mass (μg).

Separate calibration curves were made for each of the ROI methods (5*SDavg and 0.5*Max). The 5 fiducial samples imaged individually were used to create external calibration curves (Figure 1b). Correspondingly, internal calibration curves were prepared using 3 fiducial samples imaged along with Samples 1 or 2 (Figure 1c) for a total of 12 lines per user (triplicate samples x 2 test samples x 2 ROI analysis methods). The slope from the calibration curves was used to estimate iron mass from test samples, which was calculated from the MPI signal measured for each test sample:

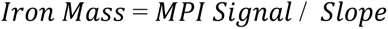

Each user estimated iron mass from Sample 1 and Sample 2 using internal or external calibration, with ROI thresholds of 5*SDavg or 0.5*Max (n = 3, total 24 measurements per user).

### Statistical Analysis

For calibration lines, the relationship between iron mass and MPI signal was determined with a simple linear regression with the y-intercept set to zero. Ordinary two-way ANOVA with Šídák’s multiple comparisons test was performed to look for significant differences between maximum signal values obtained at RRI and UF and again to compare calibration slopes at each institute.

Ordinary one-way ANOVA with Tukey’s multiple comparisons tests were performed to look for inter-user differences when obtaining measurements of iron mass with either calibration method (external or internal) and ROI segmentation method (5*SDavg or 0.5*Max). In this ANOVA analysis, lower p-values and higher F values equate to stronger differences amongst the data. Data from six users was pooled by test sample and ROI segmentation method, then student’s paired t-tests were conducted to compare iron mass estimated using external versus internal calibration. Student’s paired t-tests were also performed when comparing iron measurements made on images exchanged and analyzed between users. In all statistical tests, a p-value exceeding 0.05 was considered a statistically significant result.

### MPI Relaxometry

One user from each lab performed relaxometry for shared bottles of Ferucarbotran (Meito-Sangyo Co.), Vivotrax+ (Magnetic Insight Inc.), and Synomag-D (Micromod) to compare signal per Fe mass of the two MPI systems. The Fe content of the stock of each tracer was determined using the 1,10-phenanthroline assay [26]. A sample of each agent was prepared, with an iron mass that resulted in maximum signal intensity of ∼2 a.u.. This preparation was chosen to avoid signal saturation of the receive coil which occurs at 4-5 a.u. Samples were individually scanned using the RELAX™ module equipped on the MOMENTUM™ imager (Magnetic Insight Inc.) to produce a point spread function (PSF) for each sample. PSFs were normalized by the amount of iron scanned to produce a curve of signal per μg Fe. The maximum signal was determined in Excel (Microsoft). Additional relaxometry scans were performed on several volumes (2, 5, and 10 μL) of Vivotrax+, Ferucarbotran, and Synomag-D on each MPI system.

## Results

### Formation of Calibration Curves

Examples of external calibration and internal calibration curves are shown in Figure 1b and 1c, respectively. Figure 1b shows MPI images of 5 calibration samples used for external calibration which are arranged in order of increasing mass, from 2.5 to 40 μg. Samples with more SPION mass produced higher MPI signals (a.u.) in a linear fashion. Figure 1c depicts the internal calibration method, encompassing a test sample and three calibration samples containing 2.5, 5 and 10 μg Fe in a single image. There are two calibration curves, corresponding to application of two ROI segmentation methods: 5*SDavg and 0.5*Max. Each of these segmentation methods provided a linear relationship of signal to mass.

### Inter-user Variability and Accuracy of Iron Estimations

Overall, the average and standard deviation of user’s estimates of SPION mass in test Samples 1 and 2 was 10.65 ± 2.06 *μ* g (ground truth = 10 *μ*g). The estimation of SPION mass was significantly different between institutions for both samples, with Sample 1 being measured at 9.11 ± 0.79 *μ*g at RRI and at UF was 9.87 ± 1.37 *μ*g (p = 0.0052) and Sample 2 being measured at 10.82 ± 1.10 *μ*g at RRI and at UF was 12.93 ± 2.36 *μ*g (p < 0.0001). There were a total of 144 user measurements (6 users x triplicate images x 2 test sample volumes x 2 calibration methods x 2 ROI methods). The maximum estimated value was 17.95 *μ*g and this value was obtained using 0.5*max segmentation and quantification of Sample 2 (larger volume). The minimum estimated value was 7.4 *μ*g and this value was obtained using 5*SDavg segmentation of Sample 1 (smaller volume). All user’s estimates are depicted in Figure 2 and Supplementary Table 1. Overall, a greater number of significant differences were seen between users across institutions (34/45 = 76%) compared to users within an institution (11/45 = 24%). ANOVA demonstrates that the greatest inter-user differences occur when using 0.5*Max segmentation and external calibration to quantify Sample 2 (F = 22.33, p < 0.0001). The least user differences were seen with 5*SDavg segmentations and internal calibration to quantify Sample 1 (F = 4.82, p = 0.012). However, there is overall no consensus on whether external or internal calibration curves result in greater variability in measurements across ROI segmentation methods and the volume of the test sample. When comparing ROI segmentation methods, in general, more user variability (higher ANOVA F-values) was seen with 0.5*Max thresholds compared to 5*SDavg thresholds. The exception was when using external calibration for quantifying Sample 1 (same volume as calibration samples), for which both ROI segmentation methods provided similar user differences (Figure 2.a.i. show similar ANOVA F-values).

**Figure 2:**
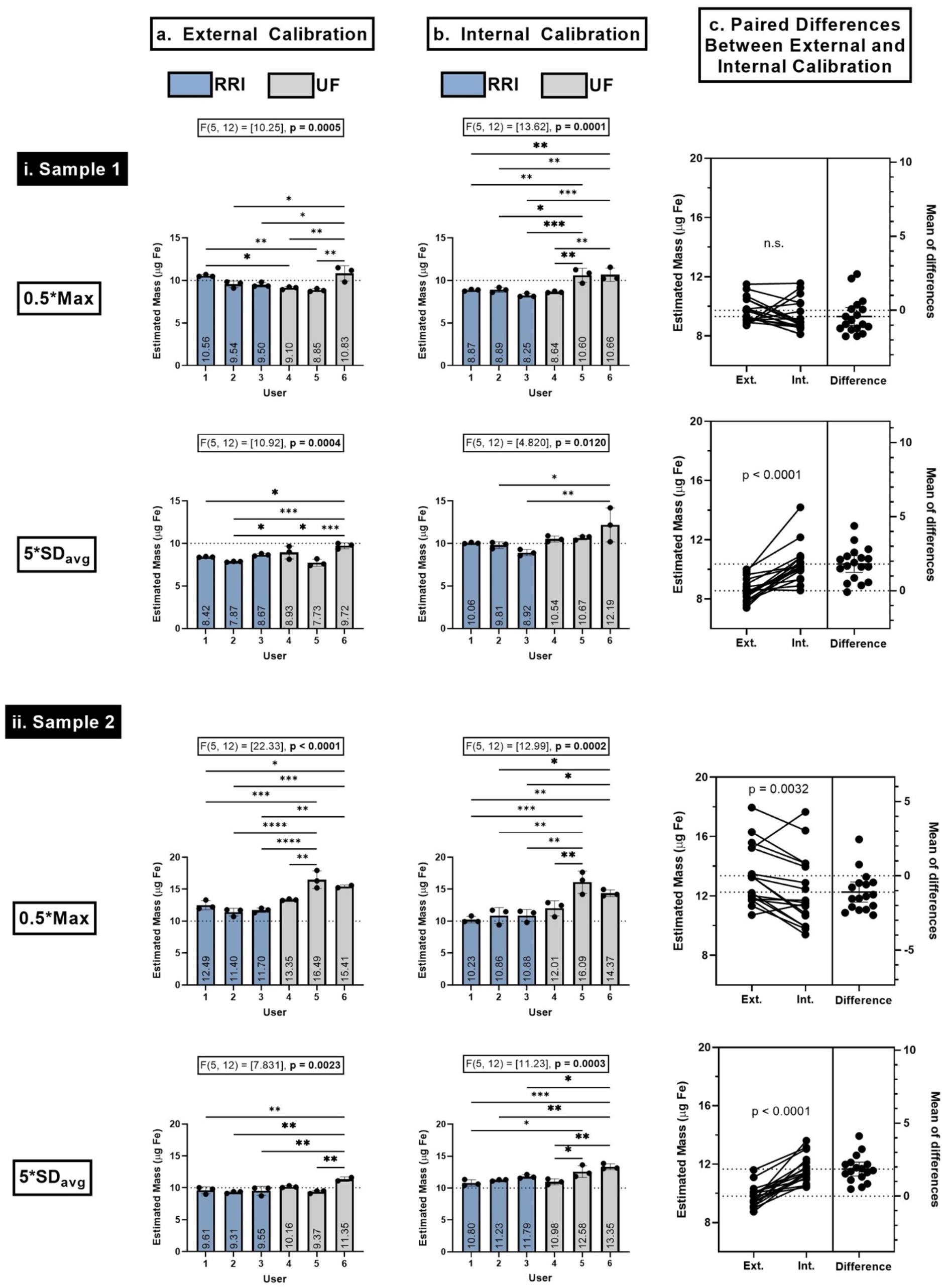
Estimation of iron mass from (i) Sample 1 and (ii) Sample 2, when using (a) external fiducials or (b) internal fiducials for calibration. These measurements were repeated for both ROI segmentation methods: at 0.5*Max (shown above) or 5*SDavg (shown below). Triplicate measurements from all Users (# 1-6) are shown, where Users 1-3 are form RRI (shown in blue) and Users 4-6 are from UF (shown in grey). The average measurements (μg) are recorded in each box. The ground truth is indicated by the dotted line at 10 μg. Statistically significant differences between user’s measurements are displayed, as determined by ANOVA (* p < 0.05, ** p < 0.01, *** p < 0.001, **** p < 0.0001). (c) The statistical difference between each user’s measurements of iron mass (μg) using external versus internal calibration methods were assessed (6 users x triplicates = 18 values). Student’s paired t-tests were performed to determine statistical significance (* p < 0.05). The mean of differences between external versus internal calibration methods are shown, where values closer to 0 indicate minimal differences between calibration techniques.

Data from all 6 users was pooled in Figure 2c (n = 18, 6 users x 3 measurements). The differences in iron mass quantified using external versus internal calibration curves were statistically significant when using 5*SDavg segmentations (Sample 1 and 2, p < 0.0001) and 0.5*Max segmentations (Sample 2 only, p = 0.0032). This indicates that measurements made using external and internal calibration methods can significantly differ. More important are differences from ground truth, which are shown in Figure 3 and Table 2. However, these comparisons do not reveal a consensus on which calibration method (internal or external) provides closer estimations to ground truth.

**Figure 3:**
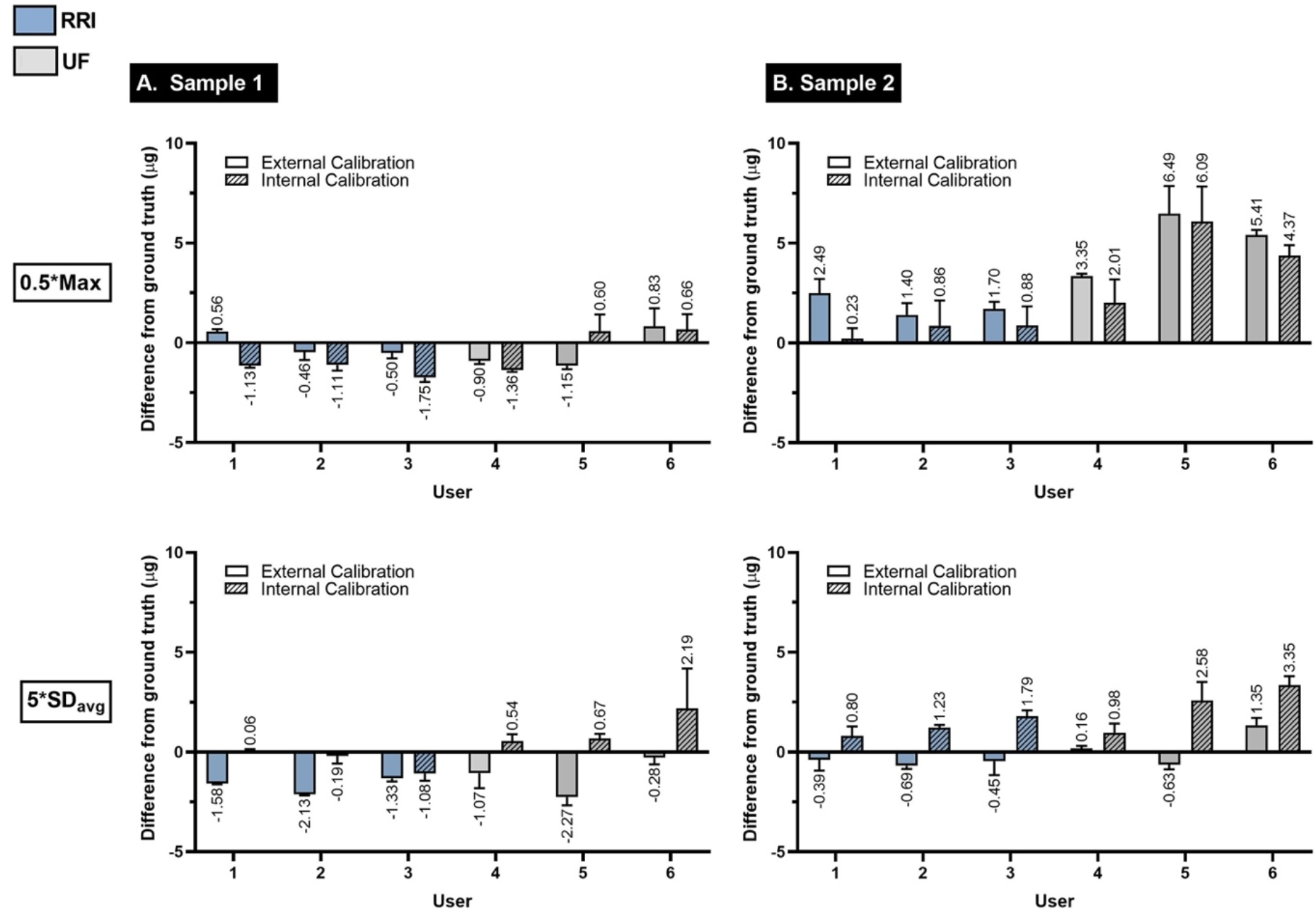
The difference (μg) between user’s estimated iron mass from ground truth (10 μg) for (A) Sample 1 and (B) Sample 2 are shown for both external (no pattern) and internal (hatch pattern) calibration techniques. The measurements made with both ROI segmentation methods are shown: 0.5*Max (above) and 5*SDavg (below). Plots show average ± standard deviation from ground truth (n = 3), where values close to 0 indicate small differences from ground truth. Users 1-3 are from RRI (blue) and user 4-6 are from UF (grey).

**Table 2:** Arithmetic averages of all user estimations of SPION mass from test Sample 1 and 2 (or their combined data) corresponding to Figure 2. The difference from ground truth values correspond to Figure 3. The value closer to ground truth is bolded for each pair of external and internal calibrations to show that no consensus can be obtained on which calibration method is more accurate.

|  |  | Average Estimated Iron mass (ug) (n = 18) |  |  | Difference from ground truth (ug) |  |
| --- | --- | --- | --- | --- | --- | --- |
| Sample | ROI | External | Internal | t-test significance | External | Internal |
| 1 | 0.5*max | 9.73 | 9.32 | No | **-0.27** | -0.68 |
| 1 | 5*SDavg | 8.56 | 10.36 | p < 0.0001 | -1.44 | **0.36** |
| 2 | 0.5*max | 13.36 | 12.41 | p = 0.032 | 3.36 | **2.41** |
| 2 | 5*SDavg | 9.81 | 11.79 | p < 0.0001 | **-0.19** | 1.79 |
| Average | 0.5*max | 11.49 | 10.86 | p = 0.016 | 1.49 | **0.86** |
| Average | 5*SDavg | 9.16 | 11.08 | p < 0.0001 | **-0.84** | 1.08 |

### MPI Raw Signal Differs Between Institutions

Next, image characteristics for images acquired at RRI and UF were compared. Histograms of voxel values, including the minimum, maximum, mean, median, and standard deviation values of the voxels were obtained (Figure 4 a,b and Supplementary Figure 1). Qualitatively, the distribution of pixel values for each sample and at each institution are similar. For example, Sample 2 (which is more dilute than Sample 1) has a wider distribution of pixels and lower intensity compared to Sample 1 at both institutions. However, samples measured at RRI had consistently higher maximum signal values compared to those measured at UF. For example, comparing images of a representative 40 μg fiducial, the RRI system produced a maximum of 3870.3 a.u. (Figure 4a,i) while UF had a maximum of 1558.8 a.u. (Figure 4b,i). As shown in Figure 4c, the average maximum signal for the scans from 3 users at RRI were statistically higher in magnitude than 3 users at UF, in all samples except for the one with a mass of 2.5 μg. Ratios for maximum signals obtained from RRI and UF are summarized in Supplementary Table 2. The average of these ratios was 3.46 ± 0.42 (RRI:UF). Interestingly, for the sample with lowest iron mass (2.5 μg), this ratio increased to 4.15 but was not statistically significant between institutions. A similar relationship is observed with calibration slopes, where RRI users obtained a higher slope than UF users and this difference was statistically significant (p < 0.0001) (Figure 4d,e). On average, the slope ratio between institutions is 3.37 ± 0.36. Importantly, slopes obtained by internal or external calibration within each institution were consistent.

**Figure 4:**
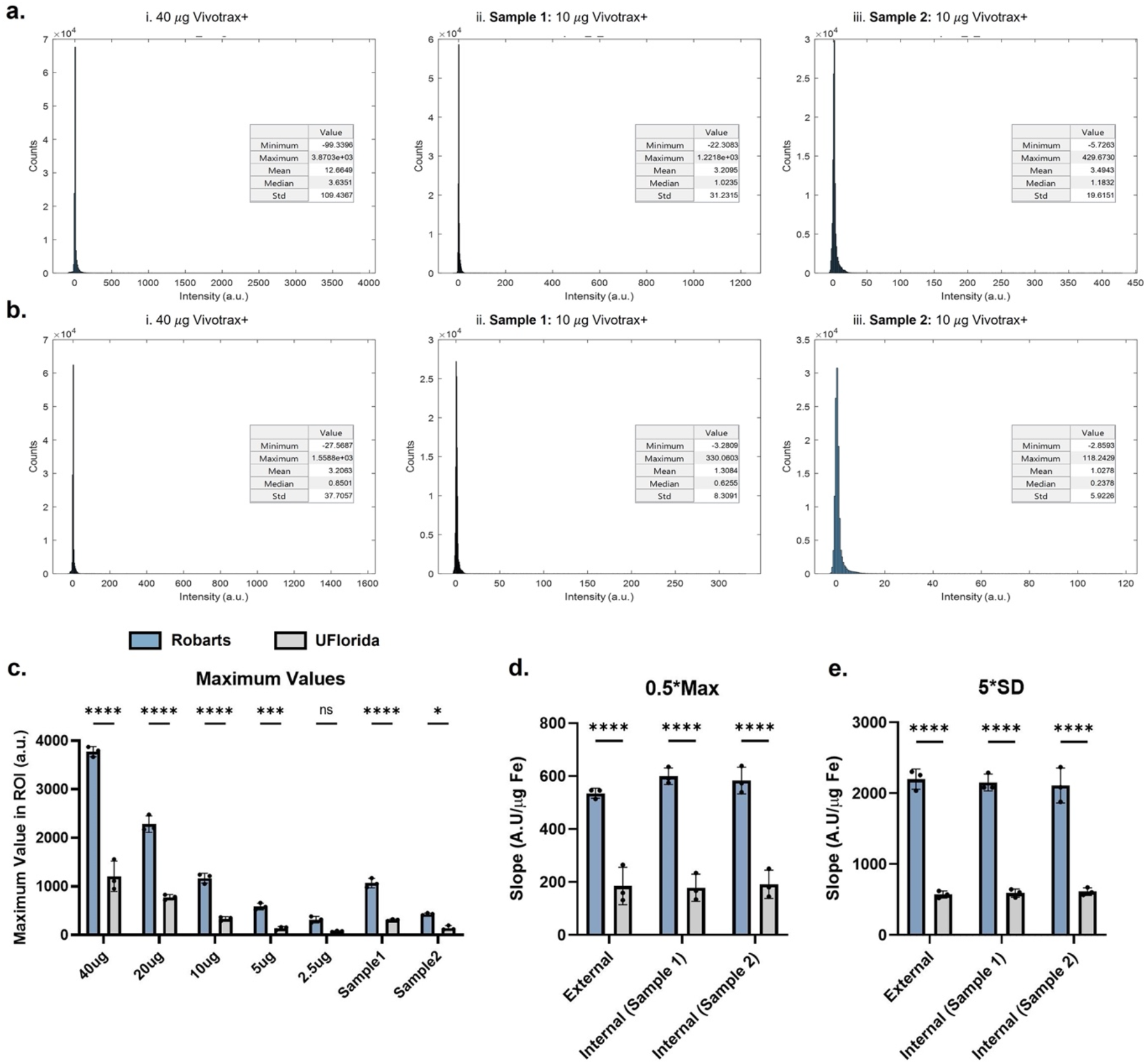
(a,b) Representative histograms show the distribution of MPI pixel values for images of i. 40 *μ*g, ii. Sample 1, and iii. Sample 2, where (a) corresponds to images acquired at RRI and (b) corresponds to images acquired at UF. The minimum, maximum, mean, median, and standard deviation (Std) of image values are reported in arbitrary units (a.u.). (c) The maximum signal from all 6 users are reported and significant differences are found between images acquired at each institution. Similarly, the slopes of the external and internal calibration curves significantly differ between institutes, for both ROI methods defined by a segmentation threshold of (d) 0.5*Max or (e) 5*SDavg. a.u. = arbitrary units, n.s. = non-significant, * p < 0.05, ** p < 0.01, *** p < 0.001, **** p < 0.0001.

### Relaxometry Results Differ Between Institutions

To further elucidate the differences of signal to SPION mass between imagers, relaxometry scans for Ferucarbotran and Vivotrax+ were performed and are shown in Figure 5. These scans show differences in maximum signal, with the ratio of maximum signals for RRI and UF was 1.322 for Ferucarbotran and 1.720 for Vivotrax+. A third SPION, Synomag-D was compared and gave a ratio between institutions of 1.566 (RRI:UF) (Supplementary Figure 2d). For these 3 SPIONs, several volumes were compared at each institute (Supplementary Figure 2 a-c). For all volumes of these SPIONs, the system at RRI produced higher relaxometry maximum signal compared to the system at UF (ratio > 1) except for measurements which saturated the receive coil (signal exceeding 4-5 a.u.). These ratios are reported in Supplementary Table 3. For all SPIONs, as iron mass in the system increased, the ratio of maximum signals from the two imagers reduced. For example, the Vivotrax+ sample shows the system at RRI has 1.582 more signal/mass than the system at UF at 2μL, 1.468 more signal/mass at 5μL, and 1.028 more signal/mass at 10μL. This is due to saturation of the detector of RRI at lower masses of SPION compared to UF. This inconsistency suggests that the relationship between signal and mass between institutions is dependent on pickup coil signal saturation.

**Figure 5:**
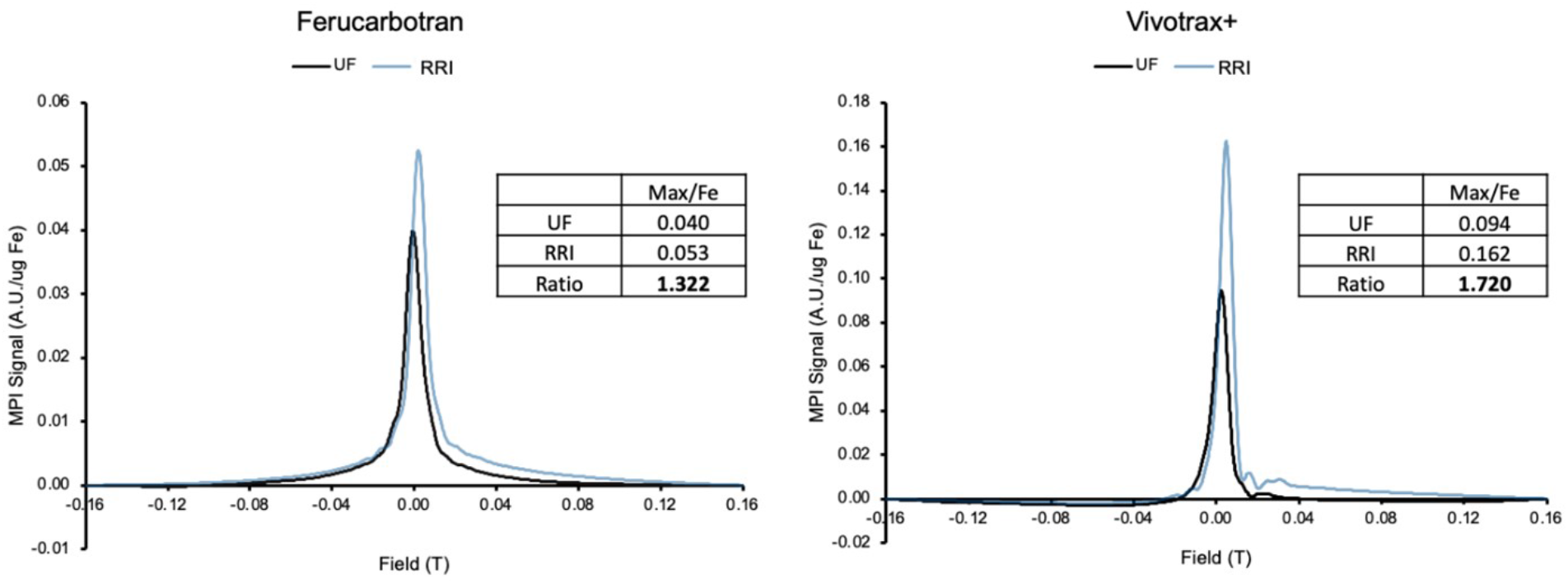
MPI relaxometry for Ferucarbotran and Vivotrax+, comparing signal strengths measured at RRI (blue) and UF (black). The ratio of peak values were obtained (RRI:UF). Optimal amounts of iron were chosen so that the raw output had maximum intensity ∼ 2 a.u. in effort to avoid saturating the detector. Relaxometry signal is normalized by the amount of iron imaged (arbitrary units/*μ*g iron).

### ROI Segmentation Approach Impacts Findings

Visual examples of ROI segmentation strategies are outlined in Figure 6 for both software programs, 3D Slicer (left column) and Horos (right column), on the same set of images obtained by User 1. Figure 6a shows the segmentations defined by the 5*SDavg approach. This segmentation commonly provided a single ROI where a manual snip was necessary to separate signal from multiple sources and form distinct ROIs. To address possible variability owing to software and these manual snips, each image was analyzed by two users (one using each software) and compared using paired t-tests, as shown in Supplementary Figure 3. This comparison shows that there was only one instance in twenty-four paired t-tests that was statistically significant. The instance corresponded to a test sample that was quantified using internal calibration and 5*SDavg segmentation method, which required four arbitrary segmentation snips to segregate the ROIs. All other comparisons showed no significant difference in quantification estimates between users and software.

**Figure 6:**
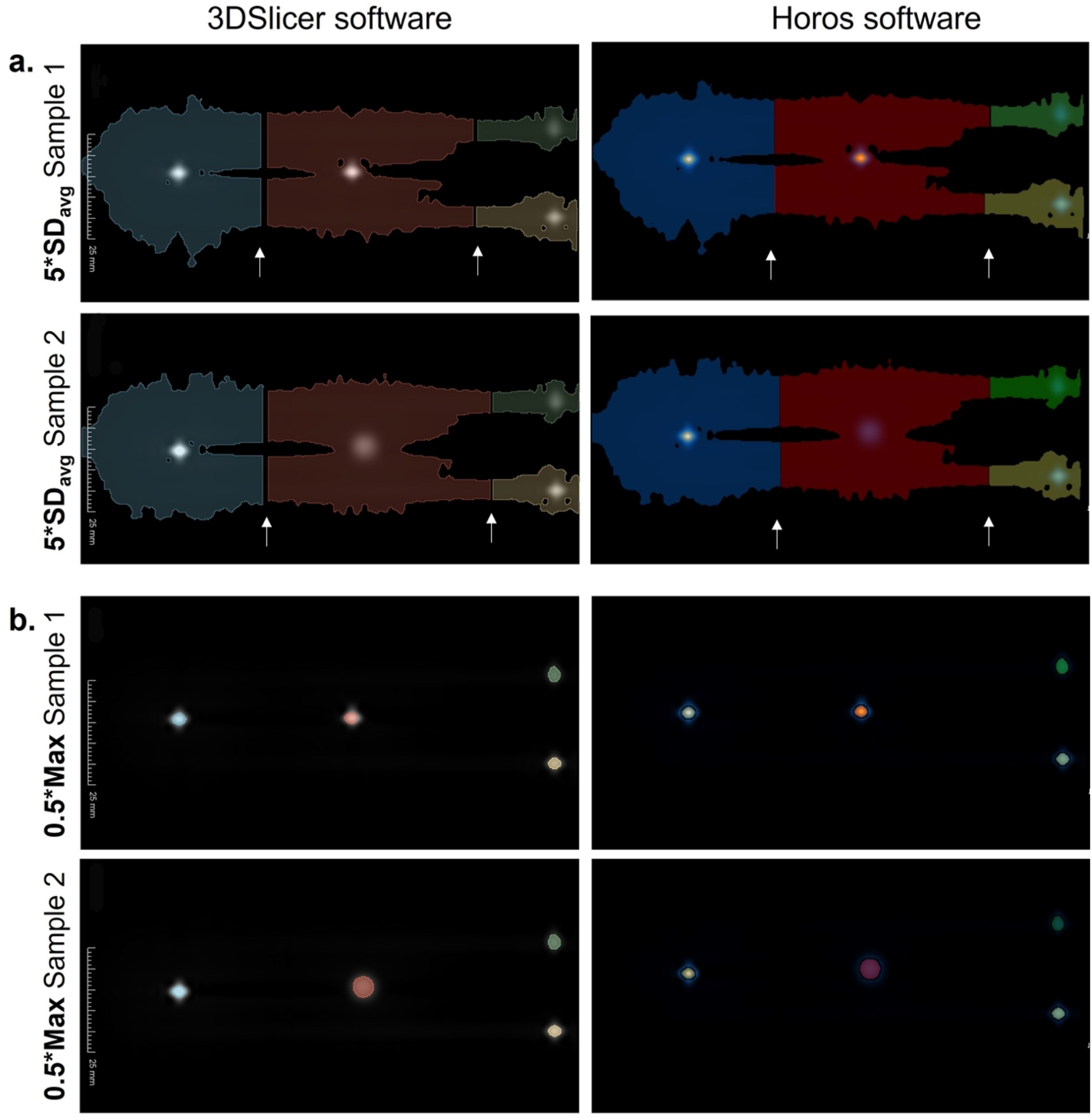
Comparison of ROIs drawn in 3D Slicer or Horos Software for images with multiple signals, using segmentations at (a) 5*SDavg and (b) 0.5*Max. The ROI for the 10 *μ*g calibration sample (left) is blue, the 5 *μ*g calibration sample (bottom, right) is yellow, and the 2.5 *μ*g calibrations ample (upper, right) is shown in green. The center ROI (red) corresponds to either Sample 1 or Sample 2, as indicated. White arrows indicate where manual separations are applied to the ROI.

## Discussion

### Scaffolds Provide Replicable Positioning

To compare imagers and prepare replicable scans, 3D printed scaffolds were designed and shared, providing consistent placement of each source of signal. These scaffolds allowed users to place signal sources in the same location in the field of view for direct comparison. This specific placement of signal sources in the field of view is important to recreate as accurately as possible, as the position of the signal source in the field of view has been shown to affect MPI scan results, as shown by Graeser, et al. [31,32].

### Calibration Method Does Not Impact Quantification Results

When comparing the two calibration methods (internal vs. external calibration), each showed similar inter-user variability and quantification accuracy across both segmentation methods and volume of sample (Figure 2). The difference in estimated iron quantification values using external and internal calibrations were statistically significant (Figure 2c and Table 2) however it cannot be said which provides more accurate results.

In comparing the two calibration methods (internal vs. external calibration), we recognize that each did not contain the same number of samples. The external calibration method contained 5 fiducials (range 2.5 – 40 μg SPION) whereas the internal calibration method contained 3 fiducials (range 2.5 – 10 μg SPION). In general, it would be expected that more calibration samples lead to overall higher accuracy in defining the calibration curve slope and a more robust relationship of signal to mass, which we would expect leads to an improved estimation of unknown mass. Nevertheless, in this study external calibration (5 samples) and internal calibration (3 samples) provided equally accurate results.

One advantage of external calibration is the flexibility to image as many samples as desired in individual images. This is a limitation for internal calibration due to practical limitations since it is challenging to separately resolve and quantify numerous signals within the same field of view. To effectively separate this signal, there must be sufficient spacing between sources to allow signal to decay toward noise and allow simple delineation of signal as performed in this study. If calibration samples produce high signals, it is possible for them to obscure MPI signals of lower intensity in the field of view, which may be relevant. If too many sources are present in the field of view, differentiation of signal sources will be convoluted and may lead to inaccuracies in the assignment of signal to source. An advantage of internal calibration method is that it reduces the total number of scans.

### Different Imagers Detect the Same Particles Differently

This study introduced a comparison between two imagers directly and quantitatively, including comparisons of maximum voxel intensities, relationships of MPI signal to SPION mass, and relaxometry measurements. In all comparisons, the imager at RRI consistently produced higher intensity signals compared to the imager at UF. When observing results for the maximum voxel intensities, RRI values were 3.46 higher than UF. Similarly, calibration slopes (a measure of signal per SPION mass) were 3.37 steeper at RRI compared to UF.

The estimation of SPION mass was significantly different between institutions for both samples, with Sample 1 being measured at 9.11 ± 0.79 *μ*g at RRI and at UF was 9.87 ± 1.37 *μ*g (p = 0.0052) and Sample 2 being measured at 10.82 ± 1.10 *μ*g at RRI and at UF was 12.93 ± 2.36 *μ*g (p < 0.0001). These differences can be attributed to different imagers and user variability, since all other aspects of the experiment were closely controlled. Results indicated that there were more differences between users across institutions (34/45) compared to users using the same imager (11/45). This suggests that the use of different imagers had a stronger influence on the estimation compared to the differences arising from user error. Furthermore, when user’s exchanged images and reanalyzed the data, there was only one instance with statistical difference. This supports the notion that differences in user’s analysis only contribute minimally to differences in SPION estimation.

Lastly, relaxometry scans of Vivotrax+, Ferucarbotran, and Synomag-D yielded consistently higher amplitude per unit mass at RRI compared to UF, with the ratio between the peak signal at the two imagers being dependent on the tracer (RRI imager 1.3 – 1.6 times higher than UF). This ratio is smaller than observed when comparing imaging signals (3.46) and calibration slopes (3.37) but shows the same trend. Raw signal intensity scaled linearly with tracer mass (controlled through stock volume used) until signal saturation occurred (5 arbitrary units). For the imager at RRI, saturation occurred at a lower amount of SPION compared to the UF imager, therefore, the ratio of relaxometry signals at institutions converge at higher concentrations of SPION. We believe these to be important observations because of the increasing interest in testing existing SPIONs and developing new SPIONs for improved MPI performance. Our observations conclusively show that merely reporting relaxometry results for a new tracer is not enough to claim improvements in signal strength or sensitivity by comparing to measurements taken for other tracers in other imagers. Furthermore, our experiments illustrate the importance of taking relaxometry measurements using samples for which the raw signal is not saturated, underscoring the need for standard data acquisition protocols.

### ROI Segmentation Approach Impacts Findings

In all instances in this study, the 5*SDavg segmentation method encompassed a larger ROI in the FOV compared to the 0.5*Max segmentations for the same image (Figure 6). Thus, more MPI signal is considered by the 5*SDavg segmentation method compared to 0.5*Max threshold, which explains why the slope of the calibration curves of data segmented at 5*SDavg are higher than that of 0.5*Max (Figure 1b,c).

Results indicate that the 0.5*Max segmentation approach led to more variability than the 5*SDavg segmentation approach (Figure 2). In the 0.5*Max method, users manually assign lower limit thresholds at 50% of the maximum signal value in that region of interest. This means that a different threshold is applied for each region of interest, unlike the 5*SDavg threshold approach which applies an identical threshold value to each signal. Since MPI scans contain discrete voxels, this thresholding will rarely be at exactly the desired value. Each user may also have chosen a different number of decimal places for their threshold values. Additionally, the threshold values used for 5*SDavg are much lower than 0.5*Max, so while these factors may seem minor or trivial, the voxels in question carry a lot of weight in the calculation of MPI signal. Therefore, each voxel included or excluded will contribute to that segmentation in a more drastic manner for the 0.5*Max method than the 5*SDavg method. These factors could contribute to the observed higher inter-user variability with 0.5*Max segmentation. However, the 5*SDavg method uses arbitrary ROI segmentations in cases where there are multiple ROI in the same image. We anticipated that this arbitrary segmentation would lead to large differences between users and iron estimation results, but instead we demonstrate that two users analyzing the same images retrieved similar quantification results. We believe that because the arbitrary segment splices are at values that are low compared to the signal source values, the assignment of those voxels in question to either ROI do not make a significant difference in the total MPI signal, thus not making a significant impact on the iron estimation.

The accuracy of SPION quantification can be influenced by the choice of ROI selection method. While each segmentation method provided similar output values for sample 1 (which has the same volume as calibration samples), 0.5*Max tends to overestimate SPION mass in the more dilute Sample 2. Evidence of this is shown in Figure 3B, where all user estimates exceed 10 μg (by on average 2.9 μg) for Sample 2 using 0.5*Max segmentation. Likewise, the highest estimation of iron mass in this study (17.95 μg) occurred with 0.5*Max segmentations (small ROI) and Sample 2 (larger volume). Conversely, the lowest estimation of iron mass in this study (7.4 μg) occurred with 5*SDavg segmentations (large ROIs) and Sample 1 (smaller volume). This reveals an important relationship between sample volume and ROI size. Larger ROIs are required to adequately capture the broader extents of MPI signals, and this is especially important when samples are diluted over a volume. 0.5*Max segmentations are known to lead to overestimation of iron mass from samples of larger volume by only considering the highest signals [19].

### Recommendations when Making Direct Comparisons between Institutions

Here, we provide recommendations on how to properly account for the differences in MPI imagers reported in this contribution. First, careful and consistent preparation of calibration fiducials is critical for quantification; these fiducials define the relationship of MPI signal to mass and inconsistencies here will directly impact results. Research publications should include detailed methods for their preparation, with accompanying data tables and plots of statistical analysis showing linearity and error of these calibration curves. Additionally, while this study did not observe significant differences between calibration methods (external versus internal), researchers should consider using both in all studies. External calibration carries the advantage of being able to include more samples in establishing the relationship between signal and mass, providing a more robust definition. However, internal calibration fiducials can demonstrate consistency with external calibration because they provide a ground truth in the field of view of the subject. Furthermore, these internal fiducials provide reference information useful for registration of optical and/or alternate instrument images with the MPI images. However, caution must be taken to choose an appropriate SPION concentration to include within the FOV, so that its signal does not obscure or is not obscured by other in-view signals. We also recommend that publications include detailed descriptions of the approaches to ROI segmentation, with accompanying sample images showing the corresponding ROI for important findings in the paper. With important findings and claims, we recommend that raw data associated with the publication should be readily available, allowing readers to verify and qualify the findings. This raw data provides a means to compare results and verify all claims by the authors. Importantly, relaxometry studies of new MPI tracers should report raw (a.u.) and specific (a.u./mg Fe) signal. Reporting these signals demonstrates that measurements were taken in the linear range of signal to iron mass. With all the above measurements, any publications with important SPION findings and claims should include comparisons with commercial tracers to qualify the findings against known tracer properties.

Importantly, these qualifying commercial tracer measurements should be with the same methodical setup as with the new tracer; we do not recommend using values from literature as they are imager specific, as demonstrated in this study. For this commercial tracer comparison, we recommend the use of Ferucarbotran (Meito-Sangyo) or alternatively VivoTrax+ (Magnetic Insight), at least until new commercial tracers with demonstrated consistent MPI properties become available. These recommendations will provide context and strong evidence of claims and findings in the upcoming publications, providing the reader with confidence in these claims.

## Conclusion

As MPI is continually growing and being explored in several biological contexts, the results from these studies must be qualified and verified across institutions. This study establishes that two MPI imagers provided statistically different quantification results using the same batch of particles, suggesting that there exists variation between imagers. Additionally, the choice of calibration method in this study did not significantly impact quantification; internal and external calibration methods provided similar results. However, the choice of segmentation analysis method in this study clearly impacted quantification; the method with the largest ROI segmentation provided improved quantification results and lower inter-user variability. This study highlights that results derived from MPI image analysis requires context to be comparable across the field. Several recommendations are made for future studies, including: (i.) adoption and reporting of detailed image acquisition and analysis protocols to reduce inter-user variability and enhance reproducibility; (ii.) inclusion of comparisons to commercial particles characterized under the same conditions to allow assessment of claims of improved tracer performance; (iii.) careful consideration of sample preparation when establishing the signal to SPION mass ratio within the imager; (iv.) detailed description of ROI segmentation approaches; and (v.) reporting of raw data to enable results verification by others.

**Author Contributions:** HJG, OCS, PJF, and CMR conceived the study and designed experiments. OCS, HJG, JJG, BY, SAM, and MAB carried out experiments, collected, and analyzed the data. OCS and HJG conducted statistical analysis of the data. HJG and OCS primarily wrote the manuscript. JJG aided revision and wrote some of the manuscript. All authors carefully read and edited the manuscript.

## Acknowledgements

**Acknowledgements:** Research reported in this publication was partially supported by the National Cancer Institute of the National Institutes of Health under award number R21CA263653, partially by the Gordon and Betty Moore Foundation under award number 9349, and partially by the National Institute for Biomedical Imaging and Bioengineering under award number R01EB031224. Paula J. Foster would like to acknowledge funding from the Canadian Institutes of Health Research. Paula J. Foster, Olivia C. Sehl, and Julia J. Gevaert would like to acknowledge funding from the Natural Sciences and Engineering Research Council of Canada.

**Conflict of Interest:** The authors indicate no potential conflicts of interest.

---

**Supplementary Figure 1:**
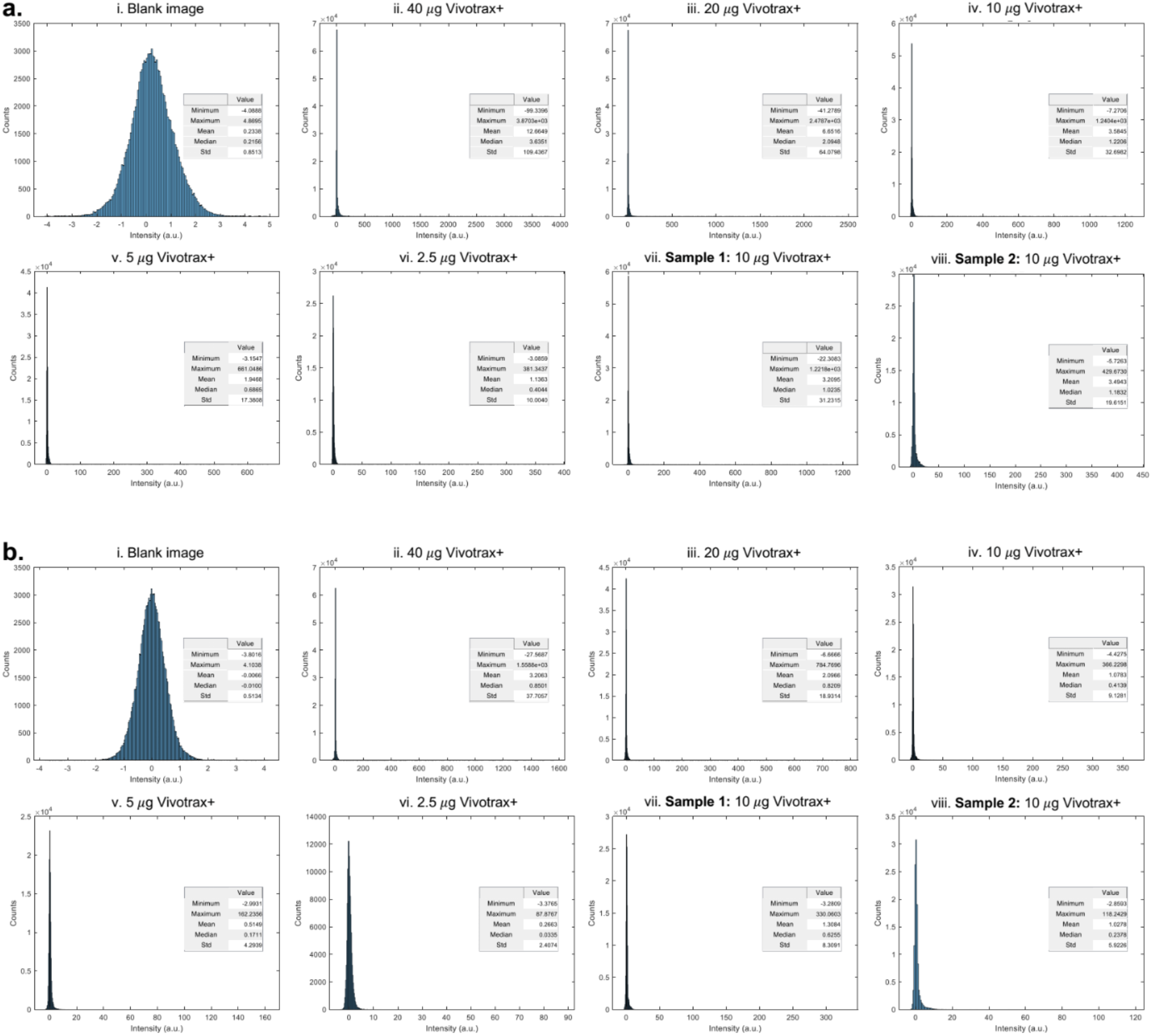
Representative histograms from images of individual samples acquired at **(a)** RRI and **(b)** UF. The pixel distribution differs for (i) blank images with no iron sample, images with Vivotrax+ calibration samples containing (ii) 40 *μ*g, (iii) 20 *μ*g, (iv) 10 *μ*g, (v) 5 *μ*g, (vi) 2.5 *μ*g and images of (vii) Sample 1, and (viii) Sample 2. The minimum, maximum, mean, median, and standard deviation (Std) of image values are reported in arbitrary units (a.u.).

**Supplementary Figure 2:**
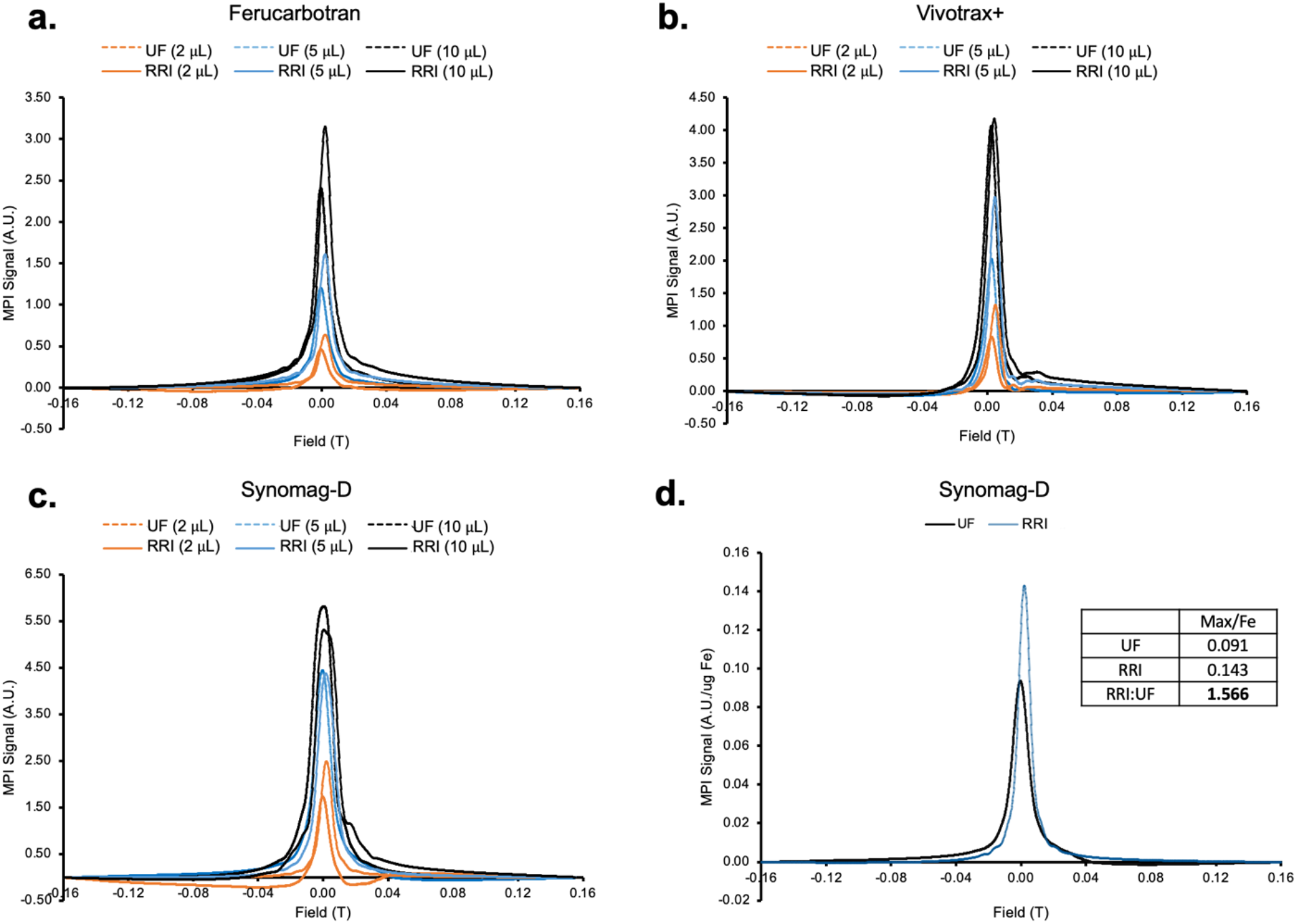
Fixed volumes (2 *μ*L, 5 *μ*L, and 10 *μ*L) from the same bottle of (a) Ferucarbotran, (b) Vivotrax+, and (c) Synomag-D were imaged at RRI (solid) and UF (dashed). (d) Optimal amounts of Synomag-D are scanned and curves normalized by iron mass to provide a sensitivity comparison.

**Supplementary Figure 3:**
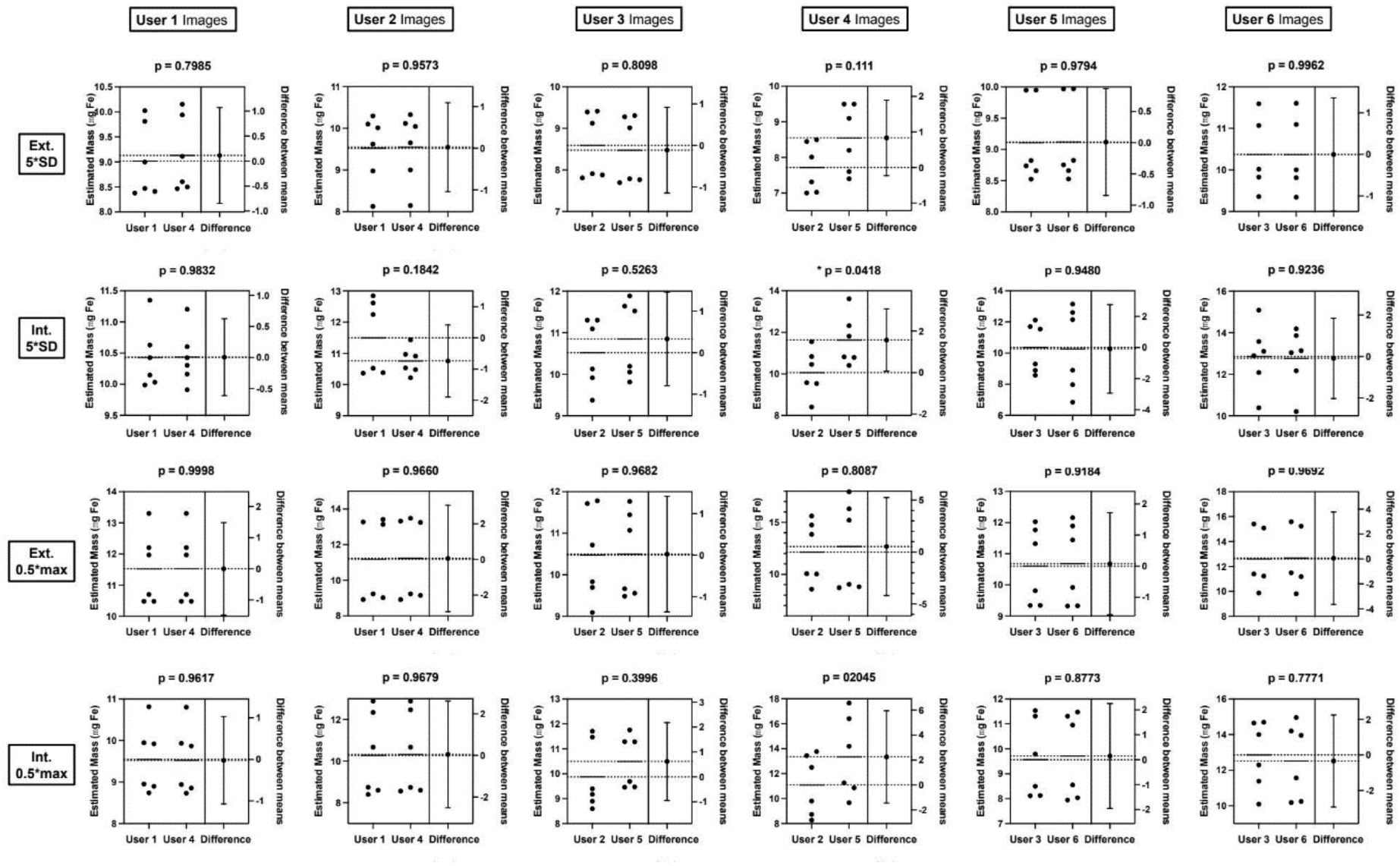
Each image was analysed by a pair of users, each belonging to a different institute (User 1-3 RRI, User 4-6 UF). Users from RRI used Horos software whereas UF used 3D slicer software. Analysis was performed using internal and external calibration using both ROI segmentation methods (5*SD_avg_ or 0.5*Max). Paired t-tests were performed between paired user’s measurements to assess for differences is image analysis (* p < 0.05).

**Supplementary Table 1:** Differences in the estimated iron mass from ground truth (μg) for each user in both segmentation (0.5*Max or 5*SD_avg_) and both calibration methods (external or internal). Values are reported as mean ± standard deviation (n = 3 per user). Institution averages are arithmetic means of the average results from the users at that institution in that mode while the provided standard deviation is the square root of the average variance from the users at that institution in that mode.

| User | Sample 1 |  |  |  |
| --- | --- | --- | --- | --- |
|  | 0.5*Max |  | 5*SD<sub>avg</sub> |  |
|  | External | Internal | External | Internal |
| 1 | 0.56 ± 0.13 | -1.13 ± 0.11 | -1.58 ± 0.05 | 0.06 ± 0.08 |
| 2 | -0.46 ± 0.39 | -1.11 ± 0.29 | -2.13 ± 0.05 | -0.19 ± 0.39 |
| 3 | -0.50 ± 0.27 | -1.75 ± 0.22 | -1.33 ± 0.15 | -1.08 ± 0.37 |
| RRI<br>Average | -0.13±0.28 | -1.33±0.22 | -1.68±0.10 | -0.40±0.31 |
| 4 | -0.90 ± 0.17 | -1.36 ± 0.09 | -1.07 ± 0.75 | 0.54 ± 0.35 |
| 5 | -1.15 ± 0.17 | 0.60 ± 0.83 | -2.27 ± 0.42 | 0.67 ± 0.23 |
| 6 | 0.83 ± 0.90 | 0.66 ± 0.78 | -0.28 ± 0.34 | 2.19 ± 1.99 |
| UF<br>Average | -0.41±0.54 | -0.03±0.66 | -1.21±0.53 | 1.13±1.17 |

| User | Sample 2 |  |  |  |
| --- | --- | --- | --- | --- |
|  | 0.5*Maximum |  | 5*SD<sub>avg</sub> |  |
|  | External | Internal | External | Internal |
| 1 | 2.49 ± 0.71 | 0.23 ± 0.51 | -0.39 ± 0.54 | 0.80 ± 0.48 |
| 2 | 1.40 ± 0.60 | 0.86 ± 1.27 | -0.69 ± 0.16 | 1.23 ± 0.12 |
| 3 | 1.70 ± 0.36 | 0.88 ± 0.95 | -0.45 ± 0.70 | 1.79 ± 0.30 |
| RRI<br>Average | 1.86±0.58 | 0.66±0.96 | -0.51±0.52 | 1.27±0.33 |
| 4 | 3.35 ± 0.12 | 2.01 ± 1.18 | 0.16 ± 0.15 | 0.98 ± 0.45 |
| 5 | 6.49 ± 1.38 | 6.09 ± 1.75 | -0.63 ± 0.23 | 2.58 ± 0.93 |
| 6 | 5.41 ± 0.25 | 4.37 ± 0.52 | 1.35 ± 0.36 | 3.35 ± 0.45 |
| UF<br>Average | 5.08±0.81 | 4.16±1.26 | 0.29±0.26 | 2.30±0.65 |

**Supplementary Table 2:** Maximum signals (a.u.) from MPI scans of the calibration samples and fiducials for Vivotrax+ at RRI and UF. Users from each institution were averaged into an institutional average, and institutional average ratios are reported.

| External Calibration Maximum Signal Values (a.u.) |  |  |  |  |  |  |  |
| --- | --- | --- | --- | --- | --- | --- | --- |
|  | Robarts |  |  | UF |  |  | RRI:UF |
|  | User 1 | User 2 | User 3 | User 4 | User 5 | User 6 |  |
| 40ug | 3870.0 | 3795.0 | 3665.0 | 1558.8 | 1111.8 | 953.2 | 3.13 |
| 20ug | 2478.0 | 2172.0 | 2197.0 | 784.8 | 827.3 | 717.8 | 2.94 |
| 10ug | 1240.0 | 1053.0 | 1216.0 | 366.2 | 349.2 | 281.1 | 3.52 |
| 5ug | 660.0 | 519.0 | 577.0 | 162.2 | 98.2 | 155.8 | 4.22 |
| 2.5ug | 381.0 | 249.0 | 317.0 | 87.9 | 62.2 | 78.4 | 4.15 |
| Sample1 | 1173.0 | 1052.0 | 983.0 | 325.2 | 287.6 | 312.4 | 3.47 |
| Sample2 | 450.3 | 407.3 | 430.7 | 117.0 | 197.4 | 112.3 | 3.02 |

| Internal Calibration Maximum Signal Values (a.u.) |  |  |  |  |  |  |  |
| --- | --- | --- | --- | --- | --- | --- | --- |
|  | Robarts |  |  | UF |  |  | RRI:UF |
|  | User 1 | User 2 | User 3 | User 4 | User 5 | User 6 |  |
| 10ug (S1) | 1247.0 | 1103.0 | 1013.3 | 390.2 | 345.7 | 197.6 | 3.60 |
| 5ug (S1) | 696.7 | 552.0 | 601.0 | 170.1 | 180.1 | 181.6 | 3.48 |
| 2.5ug (S1) | 338.0 | 229.3 | 287.0 | 97.2 | 85.2 | 87.7 | 3.16 |
| Sample1 | 1152.7 | 1017.3 | 989.3 | 331.8 | 318.9 | 294.9 | 3.34 |
| 10ug (S2) | 1281.3 | 1094.0 | 932.3 | 246.1 | 329.6 | 228.6 | 4.11 |
| 5ug (S2) | 701.0 | 557.3 | 604.7 | 177.0 | 181.9 | 186.0 | 3.42 |
| 2.5ug (S2) | 342.7 | 234.3 | 308.0 | 97.6 | 83.5 | 88.7 | 3.28 |
| Sample 2 | 446.0 | 405.0 | 384.7 | 117.0 | 182.9 | 111.7 | 3.00 |

**Supplementary Table 3:**
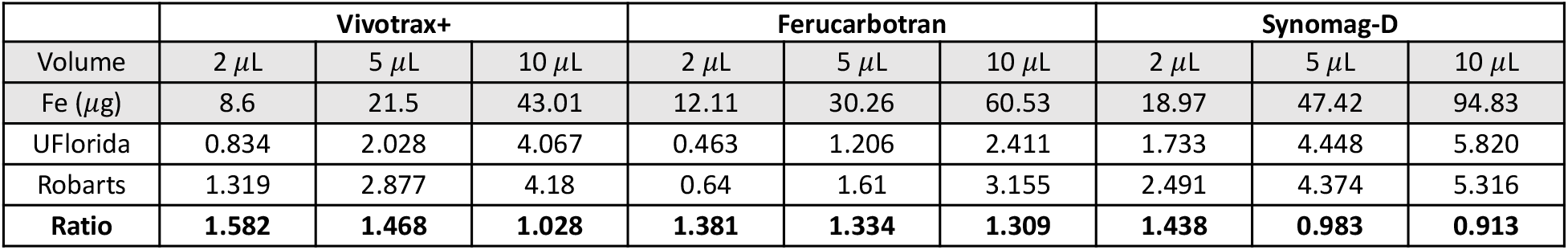
Maximum signals (a.u.) from relaxometry of fixed volumes (2 *μ*L, 5 *μ*L, and 10 *μ*L) for Vivotrax+, Ferucarbotran, and Synomag-D at RRI and UF. The signal obtained at RRI is compared to UF as a ratio (bold).

